# Proteasomal cleavage prediction: state-of-the-art and future directions

**DOI:** 10.1101/2023.07.17.549305

**Authors:** Ingo Ziegler, Bolei Ma, Bernd Bischl, Emilio Dorigatti, Benjamin Schubert

## Abstract

Epitope vaccines are a promising approach for precision treatment of pathogens, cancer, autoimmune diseases, and allergies. Effectively designing such vaccines requires accurate proteasomal cleavage prediction to ensure that the epitopes included in the vaccine trigger an immune response. The performance of proteasomal cleavage predictors has been steadily improving over the past decades owing to increasing data availability and methodological advances. In this review, we summarize the current proteasomal cleavage prediction landscape and, in light of recent progress in the field of deep learning, develop and compare a wide range of recent architectures and techniques, including long short-term memory (LSTM), transformers, and convolutional neural networks (CNN), as well as four different denoising techniques. All open-source cleavage predictors re-trained on our dataset performed within two AUC percentage points. Our comprehensive deep learning architecture benchmark improved performance by 1.7 AUC percentage points, while closed-source predictors performed considerably worse. We found that a wide range of architectures and training regimes all result in very similar performance, suggesting that the specific modeling approach employed has a limited impact on predictive performance compared to the specifics of the dataset employed. We speculate that the noise and implicit nature of data acquisition techniques used for training proteasomal cleavage prediction models and the complexity of biological processes of the antigen processing pathway are the major limiting factors. While biological complexity can be tackled by more data and, to a lesser extent, better models, noise and randomness inherently limit the maximum achievable predictive performance. All our datasets and experiments are available at https://github.com/ziegler-ingo/cleavage_benchmark.

## 1. Introduction

Epitope vaccines (EV) are a promising avenue for precision medical treatment of conditions such as autoimmune diseases and cancer, as they offer the possibility of including highly patient- and disease-specific antigens [Boniolo et al., 2021]. EVs are long polypeptides that contain the very specific parts of an antigen that are recognized by the immune system and trigger an immune response, the so-called epitopes. Proteasomal digestion plays a major role in EV efficacy [Cornet et al., 2006] (Fig. 1). By cleaving the EV incorrectly, therapeutic epitopes might get destroyed and will not be presented at the surface by the Major Histocompatibility Complex (MHC) to T cells.

**Figure 1.**
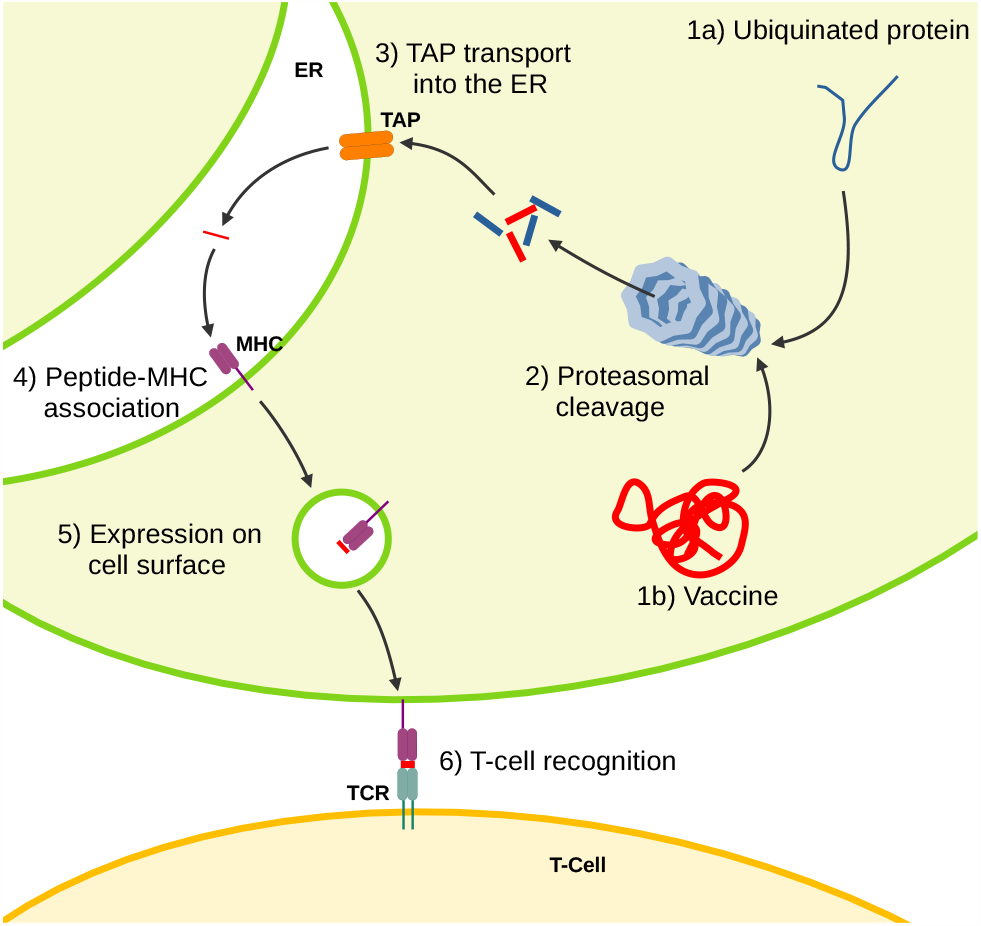
The events in the antigen processing pathway determine what is presented to T cells, thus accurate predictions are essential to enable the design of epitope vaccines. The original purpose of proteasomal cleavage (2) is to digest ubiquinated (old or misfolded) proteins into shorter peptides, but epitope vaccines (1b) undergo the same process. Some of these peptides enter the endoplasmic reticulum (ER) through the Transporter associated with Antigen Processing (TAP, 3), and a fraction of these bind to the Major Histocompatibility Complex (MHC, 4). The resulting complex is then presented on the cell surface (5) enabling immunosurveillance by T cells (6).

Several strategies have been tested to deliver EVs in the most effective way [Sette and Fikes, 2003], and it has been experimentally shown that epitope ordering in an EV affects its efficacy by determining the specificity of the immune response to antigen stimulation after vaccination [Cornet et al., 2006]. Epitope ordering affects cleavage efficiency at their terminals and thus the probability that they are correctly separated from the EV construct. Modern EV design frameworks [Dorigatti and Schubert, 2020a,b], therefore, leverage proteasomal cleavage predictors to improve epitope recovery rates, thereby improving their effectiveness at lower dosages and thus reducing the cost to patients.

In this review, we give an overview of all known proteasomal cleavage site predictors and benchmark all currentlyaccessible ones. In light of the recent developments in the field of deep learning, we additionally benchmark a wide variety of neural architectures, denoising methods, amino acid embedding methods, and training regimes, that have never been applied to proteasomal cleavage prediction before and identify parameters and architecture choices that impact performance. Based on the benchmark results and ablation study, we finally discuss the future direction of the field and give recommendations for the development of novel proteasomal cleavage predictors.

## 2. Evolution of proteasomal cleavage predictors

Proteasomal cleavage prediction efforts (Table 1) were initiated by the Prediction Algorithm for Proteasomal Cleavages (PAProC), predicting proteasomal cleavage sites of human and yeast proteasomes [Kuttler et al., 2000], which later has been made publicly available as a web-server [Nussbaum et al., 2001]. The model encodes hand-picked features such as distance information from one cut to the next as well as probabilities about neighboring amino acids and the re-sulting likelihood leading to another cut. While PAProC was only trained on *in vitro* data, Keşmir et al. [2002] also used natural MHC-I ligands and extended their initial multilayer-perceptron to an ensemble of neural networks to further improve performance [Nielsen et al., 2005a]. Cleavage prediction was integrated with the two other main processes of the antigen processing pathway, namely TAP transport and MHC binding, almost concurrently by [Tenzer et al., 2005b, ProteaSMM] and [Dönnes and Kohlbacher, 2005, PCM]. Both methods are based on position-specific scoring matrices (PSSM), where a separate coefficient is associated with each amino acid position combination and the final cleavage probability is derived linearly from such scores. Bhasin and Raghava [2005] proposed Pcleavage, a cleavage predictor based on Support Vector Machines [Cortes and Vapnik, 1995, SVM], which has been made available as a web-server. Along Pcleavage, the authors benchmarked alternative classifier systems based on nearest-neighbors, logistic regression, naive Bayes, and decision trees, but found the SVM predictor to perform best, with little performance difference between polynomial and radial basis function (RBF) kernels. Ginodi et al. [2008] approached cleavage prediction from a uniquely different perspective and tried to predict the probability that a given peptide was generated by proteasomal cleavage event, rather than the cleavage probability at each position of an antigen, with a similar approach to that of PCM.

**Table 1.** Chronological overview of existing proteasomal cleavage predictors, the modeling technique they adopted, which epitope terminals were modeled, the window size before (B) and after (A) the cleavage site, whether they used *in vitro* or mass-spectrometry (MS) training data, and the public availability of the predictor. Links to model, code and/or web server functionality, when available, are found in Appendix A.1.

| Model | Modeling Technique | Epitope Terminals | Window size |  | Training data |  | Availability |  |  |
| --- | --- | --- | --- | --- | --- | --- | --- | --- | --- |
|  |  |  | B | A | *in vitro* | MS | Model | Code | Web |
| PAPROC [Kuttler et al., 2000] | MLP | N/C | 6 | 4 | ✓ |  |  |  |  |
| NetChop [Keşmir et al., 2002] | MLP | C | 9 | 9 | ✓ | ✓ |  |  |  |
| ProteaSMM [Tenzer et al., 2005a] | Linear | C | 6 | 4 | ✓ |  | ✓ | ✓ |  |
| PCM [Dönnes and Kohlbacher, 2005] | Linear | C | 4 | 2 | ✓ |  |  |  |  |
| NetChop 3.1 [Nielsen et al., 2005a] | MLP Ens. | C | 8 | 8 | ✓ | ✓ | ✓ |  | ✓ |
| Pcleavage [Bhasin and Raghava, 2005] | SVM | C | 9 | 9 | ✓ | ✓ |  | ✓ |  |
| [Schatz et al., 2008] | Linear | N/C | 5 | 0 |  | ✓ |  |  |  |
| PCPS [Diez-Rivero et al., 2010] | N-grams | C | 6 | 6 |  | ✓ |  |  | ✓ |
| [Xie et al., 2013] | SVM | C | 14 | 14 | ✓ | ✓ |  |  |  |
| NetCleave [Amengual-Rigo and Guallar, 2021] | MLP | C | 4 | 3 |  | ✓ | ✓ | ✓ |  |
| Pepsickle [Weeder et al., 2021] | MLP | C | 8 | 8 | ✓ | ✓ | ✓ | ✓ |  |
| PUUPL [Dorigatti et al., 2022] | MLP Ens. | N/C | 6 | 4 |  | ✓ | ✓ | ✓ |  |
| This paper | BiLSTM | N/C | 6 | 4 |  | ✓ | ✓ | ✓ |  |

While most previous and following cleavage predictors focused on the C-terminal, Schatz et al. [2008] used the PSSM technique introduced previously to study N-terminal cleavage patterns and used the insights derived from their models to propose a more comprehensive modeling approach of the antigen processing pathway comprising epitope trimming besides proteasomal cleavage, TAP transport, and MHC binding. With PCPS, Diez-Rivero et al. [2010] used a model based on *n*-gram frequencies due to their success in computational linguistics and showed that their model in conjunction with MHC-I binding prediction greatly increase the discovery rate of MHC-restricted CD8 T cell epitopes. Xie et al. [2013] departed from the one-hot encoding of amino acids used in the previous methods and instead used amino acid features summarizing their physio-chemical properties [Mei et al., 2005, VHSE] via principal-component analysis [Jolliffe, 2002, PCA]. By interpreting the importance of the features used by their proposed SVM-based predictor, the authors identified proteasome specificities in terms of hydrophobicity of the amino acids in the flanking regions. NetCleave [Amengual-Rigo and Guallar, 2021] extended cleavage prediction to MHC-II and H2 ligands and returned to neural network models, while also using VHSE amino acid descriptors. Pepsickle [Weeder et al., 2021] collected a larger dataset of *in vitro* cleavage sites and showed that using predicted C-terminal cleavage probabilities helped isolating immune-responsive epitopes, thus improving EV design. Finally, Dorigatti et al. [2022] proposed to treat decoy samples as unlabeled, rather than negatives, and used specific techniques that do not require labeled negatives to learn a classifier [Bekker and Davis, 2020].

### 2.1. Data types for training proteasomal cleavage prediction models

Proteasomal cleavage sites are usually inferred implicitly from MHC ligandomics data [Purcell et al., 2019] by matching eluted ligands to their progenitor protein to recover sequence information surrounding the terminals [Keşmir et al., 2002]. This procedure, however, does not give an indication of which amino acid sequences *cannot* result in a cleavage event, since missed cleavage sites are not observed in MHC ligands, and presents a biased picture of cleavage patterns as not all proteolytically generated peptides will be presented by the MHC. Alternatively, *in vitro* collected proteasomal digested proteins are used to train cleavage predictors. These data are more comprehensive and less biased, however, low-throughput and are harder to obtain compared to MHC ligandomics. Additionally, such data do not contain any information about alternative antigen-processing pathways that lead to peptide presentation and successive antigen-processing steps. This complicates the use of predictors trained on *in vitro* data alone in pipelines aimed at identifying immunogenic peptides as it creates the necessity of modeling successive pathway events separately, as done for example by NetCTLpan [Stranzl et al., 2010].

Therefore, most modern proteasomal cleavage predictors are trained solely on MHC ligandomics data or integrate both *in vitro* and *in vivo* data.

To bypass MHC liganomdics’ inability to identify true non-cleaved sites, decoy negative samples are usually generated synthetically either by randomly shuffling the amino acids in a short window around the cleavage site or by considering artificial negative sites located around observed cleavage events [Calis et al., 2014] (Fig. 2). Even though such negative samples are not entirely reliable, the growing availability of this kind of data [Vita et al., 2018] spurred continuous development and improvement of proteasomal cleavage predictors [Keşmir et al., 2002, Kuttler et al., 2000, Dönnes and Kohlbacher, 2005, Nielsen et al., 2005b], which have been recently revised in light of new innovations in the deep learning field [Amengual-Rigo and Guallar, 2021, Dorigatti et al., 2022, Weeder et al., 2021].

**Figure 2.**
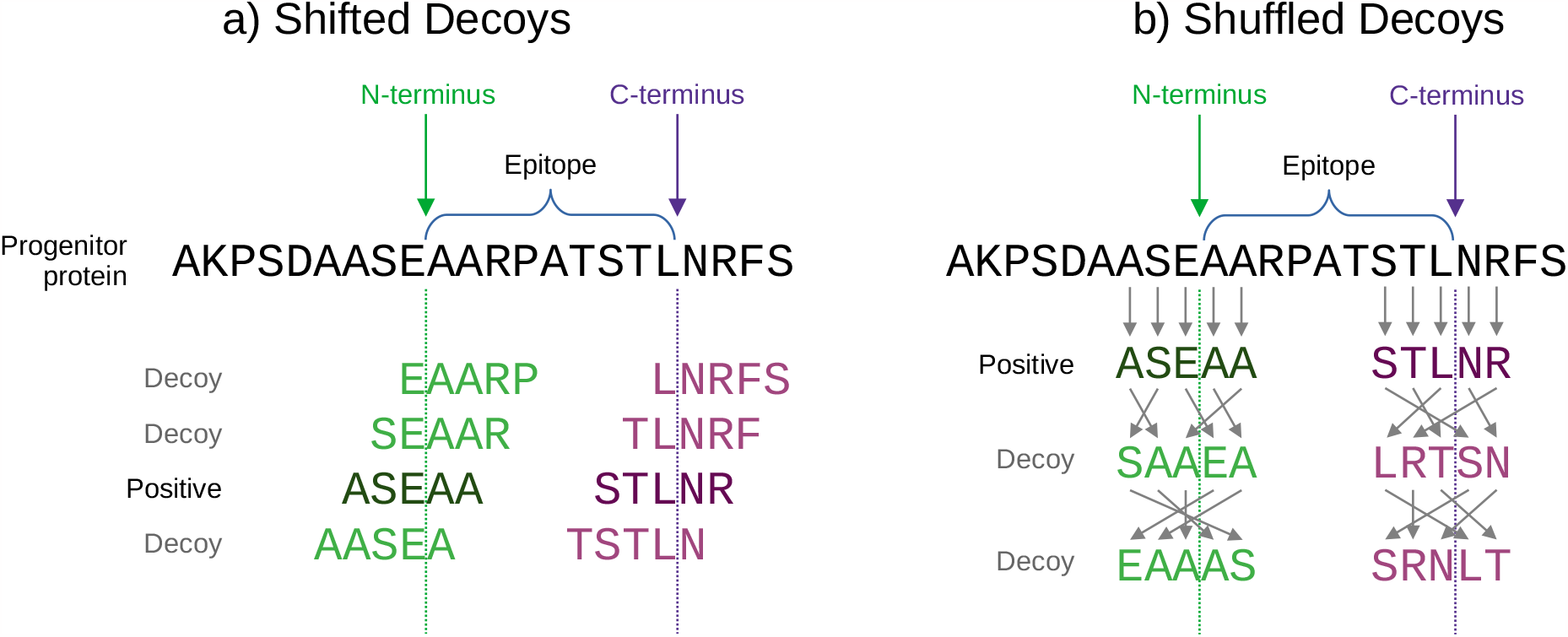
Two alternative ways of creating decoy negative samples from MHC ligandomics data. MHC-I epitopes are matched to their progenitor protein and short windows around its terminals, in the figure composed of three residues before and two after, used to construct decoys. Shifted decoys (a) are substrings of the original sequence starting earlier or after the observed cleavage site, whereas shuffled decoys (b) contain the same amino acids as the positive example but shuffled in random order. In this work, we consider shifted decoys as they are more realistic (see main text).

## 3. Benchmarking of proteasomal cleavage prediction strategies

All existing methods differ by subtle details such as the size and composition of the training data, the dimension of the window surrounding the cleavage site, and the way amino acids are encoded and presented to the model. Therefore, we performed a comprehensive benchmark of publicly available approaches on comparable datasets to compare and identify beneficial modeling approaches. Additionally, we extended this benchmark to neural architectures and modeling techniques that have not yet been used in the context of proteasomal cleavage site prediction. We considered three main axis of variation: the initial embedding of amino acids, the neural architecture of the predictor, and their training regime via noise handling and data augmentations.

### 3.1. Embedding

The concept of embeddings originally stems from natural language processing (NLP) and language models, but targeted adaptations for amino acid sequence data have already been presented [Ibtehaz and Kihara, 2021]. By associating numerical vectors to *tokens* such as characters or words, embeddings constitute the first encoding step of a model. The method employed to generate embeddings is crucial as it influences what intrinsic information a model can exploit for classification [Ibtehaz and Kihara, 2021]. We, thus, considered various embeddings in our analysis, while keeping the base architecture equal. Specifically, we analyzed the performance of a randomly initialized embedding layer that was optimized in conjunction with the whole model, as well as dedicated, pre-trained large protein language models such as Prot2Vec [Asgari and Mofrad, 2015]. Furthermore, we also employed sequence embeddings obtained by concatenating independently-trained forward and backward amino acid representations of each input [Heigold et al., 2016], as well as embeddings of the *N* most common variable-length substrings identified via byte-level byte pair encoding [Sennrich et al., 2016, BBPE] and WordPiece [Schuster and Nakajima, 2012, WP]. For BBPE, we try both *N* = 1000 and *N* = 50 000, while for WP we use *N* = 50 000.

### 3.2. Neural architectures

Different neural architectures embody different inductive biases that may or may not help a learning task. Most proteasomal cleavage methods made use of multi-layer perceptrons (Table 2), however, other types of architectures also provided high performance in sequence-based classification tasks. We, therefore, evaluated all modern types of neural architectures, from recurrent to transformer-based, convolutional neural networks, and perceptrons.

**Table 2.** AUC (in %) of several cleavage prediction methods on our dataset. We present our BiLSTM results with a window size (in parentheses) comparable to the other methods as well as with the largest window size we could use for training. ^†^ SVM, logistic regression, NetCleave, Pepsickle, and PUUPL were re-trained from scratch on our dataset, while we used the published NetChop pre-trained models and the PCPS web-server functionality to obtain their performance on our test set. We present the performance of the original PCM as well as a version re-trained on our dataset.

| Method | Dataset |  |
| --- | --- | --- |
|  | C-terminal | N-terminal |
| PCPS | 51.3 | 50.0 |
| PCM | 64.5 | 52.4 |
| NetChop 3.1 (20S) | 66.1 | 52.7 |
| NetChop 3.1 (C-term) | 81.5 | 51.0 |
| SVM<sup>†</sup> | 84.8 | 73.2 |
| PCM<sup>†</sup> | 85.3 | 75.5 |
| Logistic Regression<sup>†</sup> | 86.2 | 76.2 |
| NetCleave<sup>†</sup> | 86.9 | 76.4 |
| PUUPL<sup>†</sup> | 87.2 | 78.0 |
| Pepsickle<sup>†</sup> | 88.1 | 78.9 |
| BiLSTM (6+4) | **89.8** | **80.6** |
| BiLSTM (28+28) | **92.8** | **89.4** |

#### Recurrent

Bidirectional long short-term memory (BiL-STM) networks [Graves and Schmidhuber, 2005] are well suited for a wide range of text classification tasks, thus, as proteasomal cleavage prediction is performed on amino acid sequences in a sequence-to-one classification setting similar to text classification, we based nine of 12 model architectures around BiLSTMs. The fundamental structure for our BiLSTMs was built around the architecture proposed by [Ozols et al., 2021], in which multiple sequential BiLSTMs were followed by a hidden and an output layer. For eight of our nine BiLSTM-related experiments, we chose two sequential BiLSTMs in which sequence dimensionality was reduced by taking the maximum value of the depth-wise per-residue output of the last layer. For the hidden layer, we used the Gaussian Error Linear Units [Hendrycks and Gimpel, 2016, GELU] activation function. As the attention mechanism has also been applied to non-transformer architectures, we additionally employed a modified version of the BiLSTM with extended residual connections and an attention mechanism in order to improve its performance on protein sequence data [Liu and Gong, 2019].

#### Transformers

Besides RNNs, the attention mechanism introduced by [Vaswani et al., 2017] enabled a whole new architecture capable of processing sequences: the Transformer. These complex multi-million and up to multi-billion parameter models feature an encoder and decoder system and are frequently pre-trained on large datasets in a self-supervised manner, for example by predicting randomly masked-out tokens from an input sequence. The encoders and/or decoders of such pre-trained models are then used as the basis of new models that are fine-tuned for a variety of specific downstream tasks. We used ProtTrans’ T5-XL encoder-only model [Elnaggar et al., 2022] featuring 1.2 billion parameters and pre-trained on the UniRef dataset, as well as the ESM2 Transformer [Lin et al., 2022] in its 150 million parameter version. Additionally, we included a fine-tuning performance of ESM2 by adding a linear layer projection from its vocabulary-sized per-residue RoBERTa Language Model Head [Liu et al., 2019, Rives et al., 2021] to our binary classification target.

#### Convolutional and Perceptron

While convolutional neural networks [LeCun et al., 1998, CNN] are predominantly used in computer vision problems, the convolution filters can also be seen as a sliding *k*-mer window over the input sequence [Cavnar and Trenkle, 2021]. While CNNs have already been used in conjunction with attention for protease-specific substrates and cleavage sites [Li et al., 2019], their use for proteasomal cleavage prediction remains unexplored (Table 2). Furthermore, as most of the proposed proteasomal cleavage predictors make use of multi-layer perceptrons (Table 2), we also included a single hidden layer perceptron [Rumelhart et al., 1986] with Rectified Linear Units [Agarap, 2018] as activation function into the analysis.

### 3.3. Training

The training dataset and procedure also impacts the final performance. As described in the introduction, decoy negative samples may contain some positives. To account for negative label uncertainty, we tested multiple procedures that account for this fact. Data augmentation also proved successful to reduce overfitting and improve generalization.

#### Dataset

We used the dataset introduced in Dorigatti et al. [2022], which contains 229 163 and 222 181 N- and C-terminals cleavage sites, respectively. Each cleavage site is captured in a window comprising six amino acids to its left and four to its right and is associated with six decoy negative samples obtained by considering the three residues preceding and following it (Fig. 2), resulting in a total of 1 434 989 and 1 419 501 samples after deduplication for N- and C-terminals. As the decoy negatives are situated in close proximity to real cleavage sites and due to the probabilistic nature of proteasomal cleavage, some of the negative samples are likely to be actual, unmeasured cleavage sites, and may influence the performance of predictors trained using such data. However, we prefer this method of generating decoys compared to the popular alternative of randomly shuffling amino acids in the window as the latter results in sequences that are potentially not observed in nature, thus making the prediction problem artificially easier and overestimating the models’ performance on natural sequences. For the ablation analyses, we used the above-described dataset to investigate the impact of varying the window size around the cleavage site as well as the number of training samples used to train the predictors.

#### Noisy labels

To reduce the impact of asymmetric label noise on the performance of our classifiers, we took five recent deep learning-specific denoising approaches into consideration: a noise adaptation layer, which attempts to learn the noise distribution in the data [Goldberger and Ben-Reuven, 2017], co-teaching, where two models are trained simultaneously by deciding for the respective other network which samples from a mini-batch to use for training [Han et al., 2018], and co-teaching-plus [Yu et al., 2019], which extends co-teaching with the disagreement learning approach of decoupling [Malach and Shalev-Shwartz, 2017]. We additionally considered a joint training method with co-regularization [Wei et al., 2020, JoCoR] that, unlike co-teaching and its variations, aims at reducing the disagreement of two networks by calculating a joint loss under co-regularization and choosing small-loss samples to update the parameters of both networks simultaneously. We also tried DivideMix [Li et al., 2020a], a holistic approach originally developed for computer vision, which integrates multiple frameworks such as co-teaching and MixMatch [Berthelot et al., 2019] into one. Following the co-teaching approach, each network feeds the other network parts of the training dataset. In this case, however, the division of samples happens at the beginning of an epoch by fitting a Gaussian Mixture Model [Reynolds, 2009, GMM] and splitting the dataset at a chosen threshold. High-loss samples, as classified by the GMM, are treated as unlabeled and semi-supervised learning techniques based on MixMatch are followed. MixMatch unifies consistency regularization, where different augmented views of the same example are constrained to lead to the same prediction, with entropy regularization, which forces the classifier’s predictions to have low entropy. As MixMatch was developed for image data, we adjust it for sequence data by mixing up the embedded sequence representation [Guo et al., 2019] instead of the pixel input in the data loading process.

#### Data augmentation

For all models except external baselines as well as models using BytePair or WordPiece tokenization, we applied data augmentation directly on the input sequences to combat overfitting and improve generalization by randomly masking max (1, ⌊*𝓁*/10⌋) amino acids for a sequence of length *𝓁* as unknown [Shen et al., 2021]. The models featuring BytePair or WordPiece tokenization do not benefit from additional data augmentation because the trainable tokenization process with large vocabulary sizes might assign a single token to a whole input sequence. Consequently, the augmentation process would mask the sole token, which represents the whole sequence, as unknown.

All predictors except ESM2 fine-tuning used adaptive momentum [Kingma and Ba, 2015] as their optimization technique, whereas ESM2 fine-tuning used adaptive momentum with decoupled weight decay [Loshchilov and Hutter, 2017]. All models without denoising techniques used binary cross-entropy loss [Cox, 1958], while all denoising models calculated dedicated losses.

### 3.4. Baselines

As baseline models, we considered a logistic regression classifier with L2 regularization, which is essentially equivalent to ProteaSMM, tuned via 10-folds cross-validation, and a SVM model fitted using 15 000 samples to computational limitations, optimizing its hyperparameters via three-folds cross-validation. As for published predictors, we bench-marked against NetCleave [Amengual-Rigo and Guallar, 2021], Pepsickle [Weeder et al., 2021], and PUUPL [Dorigatti et al., 2022], retraining them on our dataset of both N- and C-terminals, as well as comparing a retrained version of PCM [Dönnes and Kohlbacher, 2005] with the published pretrained model. We also considered PCPS (Model 1) [Diez-Rivero et al., 2010], NetChop 20S and NetChop C-term [Nielsen et al., 2005a], however, we could not re-train these due to the unavailability of the source code. We could not use PAProC as the web-server [Nussbaum et al., 2001] is currently not functional, while [Xie et al., 2013] never published code nor pretrained models. Both authors did not respond to our inquiries.

### 3.5. Experimental protocol

The experimental protocol determines how we optimized and evaluated the models.

#### Hyperparameter optimization

To gather initial broad ranges of well-performing hyperparameters, we performed manual try-out runs in a grid search approach. Due to computational limitations, we divided the hyperparameter search into two priority groups: group one used Ray Tune’s [Moritz et al., 2018] implementation of the asynchronous Hyperband algorithm [Li et al., 2020b] and evaluated each configuration in a ten-folds cross-validation (CV), while for group two we chose hyperparameters according to the initial grid search results and evaluated each configuration with five-folds CV(as shown in Table 16 of Appendix A.3). We then used the best hyperparameter combination to train each architecture with all denoising methods, except for DivideMix where we only trained the overall best-performing architecture due to computational limitations. Information on the different architectures and the exact Hyperband ranges and chosen hyperparameters for all models are documented in Appendix A.2.

#### Evaluation

As previously mentioned, some negative samples may actually result in a proteasomal cleavage event *in vivo* due to the way these negative samples are generated. For this reason, traditional binary classification metrics such as accuracy, precision, recall, etc., are misleading and model evaluation should instead be based on the AUC [Menon et al., 2015]. We reserved a random 10% of each terminal dataset as test dataset used for the final evaluation of the best hyperparameters.

## 4. Results

### General results

The overall best-performing model architecture for both C- and N-terminus as measured by AUC was the BiLSTM at 89.85% (C-terminus) and 80.62% (N-terminus) without any denoising methods (Fig. 3 and Appendix A.3 and A.4). If denoising techniques were applied, the noise adaptation layer consistently performed best for both the C- and N-terminal. For the C-terminus, the noise adaptation layer showed improved performance in 5 out of 12 models. However, within these 5 models, improvements were minimal, with 4 of 5 models exhibiting less than 0.1 percentage points (pp.) enhancement. This level of improvement is negligible, particularly given that the standard deviation of the models was higher in 2 of the 4 models. In the case of the 1.2 billion BiLSTM+T5 model, the noise adaptation layer led to a more substantial improvement of 0.51 pp, resulting in an AUC of 87.21%, which is still 2.64 pp. below the 4.66 million parameter BiLSTM with 89.85 % AUC.

**Figure 3.**
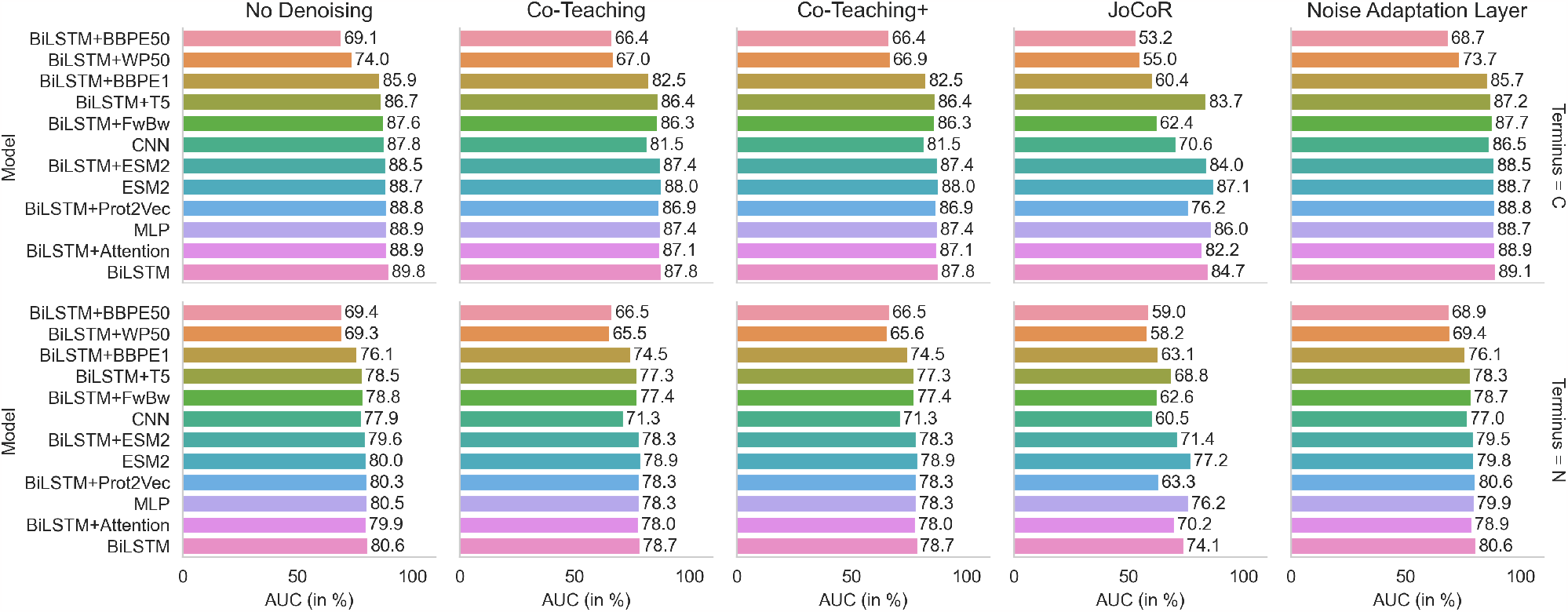
Model performances on C- and N-terminal (rows) for different denoising methods (columns).

On the other hand, for models in the N-terminus category also utilizing the noise adaptation layer, improvements were similarly marginal. The BiLSTM+Prot2Vec architecture experienced a modest increase of 0.15 pp., narrowly above its standard deviation of 0.12 pp. The BiLSTM+WP50 model exhibited an improvement of 0.07 pp., which falls below its standard deviation of 0.11 pp. In all other experiments, none of the denoising methods yielded superior performance.

### Denoising methods

In the context of experiments with other denoising methods, it is noteworthy that co-teaching-plus yielded identical outcomes in 11 out of 12 models for both the C- and N-terminals. This indicates that the decoupling-based disagreement learning approach within co-teaching-plus was not triggered except once, resulting in co-teaching-plus being equal to co-teaching. In the one remaining scenario of the C-terminus, co-teaching-plus showed slightly inferior results, trailing co-teaching by 0.14 pp. Nevertheless, this difference was still within the narrow standard deviation range of 0.15 pp. For the N-terminus, AUC marginally increased by 0.1 pp. for the case where co-teaching-plus took effect. This time, the performance gain of 0.1 pp. exceeded the model’s standard deviation range of ±0.08 pp. For both the C- and N-terminal, the BiL-STM+WP50 model activated co-teaching-plus’s disagreement learning approach.

JoCoR appeared to significantly hinder model performance in all architectures and across both terminals. For the C-terminus, the losses range from -1.62 pp. for the ESM2 model to around -25 pp. for both the BiLSTM+FwBw and BiLSTM+BBPE1 models. The N-terminus saw similar but less pronounced performance deficiencies between -2.79 pp. on the low end for the ESM2 model and up to -20.58 pp. for the BiLSTM+WP50 model. DivideMix, as the most encompassing denoising method, also reduced the BiLSTM AUC score by around 3.6 pp. in both terminals.

### Performance plateaus

While the best-performing BiL-STM consisted of 4.6 million parameters, the MLP with 30 529 parameters only lacked 0.99 pp. AUC behind. Most surprisingly, the MLP outperformed 9 out of the remaining 11 architectural configurations, achieving an AUC of 88.86%. This implies that a small, from-scratch-trained model with 30 529 parameters surpassed all the pre-trained Prot2Vec embedding models, the 148 million pre-trained and fine-tuned ESM2 transformer, the 152 million BiL-STM+ESM2 combination, and the 1.2 billion BiLSTM+T5 model, as well as all other BiLSTM and tokenizer combinations. Only the BiLSTM and BiLSTM+Attention models exhibited superior performance, suggesting that the general pre-trained Prot2Vec embeddings may have hindered the performance of the third BiLSTM model, the BiL-STM+Prot2Vec, in this specific task. Although the test set performance of the MLP, indeed, ranked 3^rd^, it is noteworthy that the performance differences to the non-trainable-tokenizer models ranged only from 0.06 pp. to 2.95 pp. This highlights more of a general performance plateau rather than the dominance of a single architecture. Models trained on the N-terminal followed a similar pattern.

### Trainable tokenizers

For the C-terminal, models including trainable tokenizers dropped to their worst-performing state compared to their fixed-vocabulary counterpart, especially when increasing the number of amino acid substring combinations that had to be learned. Whereas the BiLSTM with BBPE vocabulary size 1000 dropped 3.9 p.p. to 85.94% AUC, the same model architecture with 50 000 learned sub-string combinations was only able to reach 69.14% AUC. A similar but less severe pattern could be observed with WordPiece encodings, where the version with vocabulary of size 50 000 reached 73.95% AUC. If these models additionally featured denoising methods, the performance loss intensified up to a level of almost random-guessing with performances losses in the ranges of up to 37.65 pp. (53.20% AUC for 50 000 BBPE and JoCoR, 55.01% AUC for WP and JoCoR).

On the other hand, the performance loss from to trainable tokenizers for N-terminal cleavage sites was less severe. Although the direct comparison between the BiLSTM and BiLSTM+BBPE1 for C- and C-terminal was larger with -4.5 pp. instead of -3.9 pp., the maximum performance losses are limited at 11.2 pp. for both BiLSTM+BBPE50 and BiLSTM+WP50. When trainable tokenizers are paired with JoCoR, the performance also drops to their lowest level. However, the largest performance loss is capped at 22.45 pp. for BiLSTM+WP50 compared with the BiLSTM+BBPE50 at 37.65 pp. of the C-terminal. Replacing the embedding layer with a forward-backward representation does not benefit the classification task. Compared with the BiLSTM, not only does performance drop around 2.3 pp., the runtime also increases by a factor of six due to the necessity of increased sequential modeling of each forward-backward token representation.

### 4.1. Ablation results

To study the influence of the dataset on the predictive performance, we trained the best-performing model identified in the comprehensive benchmark, the BiLSTM, with fewer and fewer training examples as well as with different window sizes around cleavage sites. Due to the small variation in performance discussed earlier, it is likely that repeating these experiments with different methods would result in qualitatively identical conclusions.

#### Training set size

The size of the training dataset had a negligible impact on performance, with a decrease of only 1.08 and 0.98 p.p. in AUC with half as many training samples for C- and N-terminals respectively (Fig. 4 left, and Table 29 in Appendix A.5). The performance decrease when training with only 10% of the full dataset (about 22 000 cleavage sites and 140 000 decoys) was 2.7 p.p. for N-terminals and 2.06 for C-terminals. We monitored training and validation losses throughout to ensure that no overfitting occurred with the smaller training sets.

**Figure 4.**
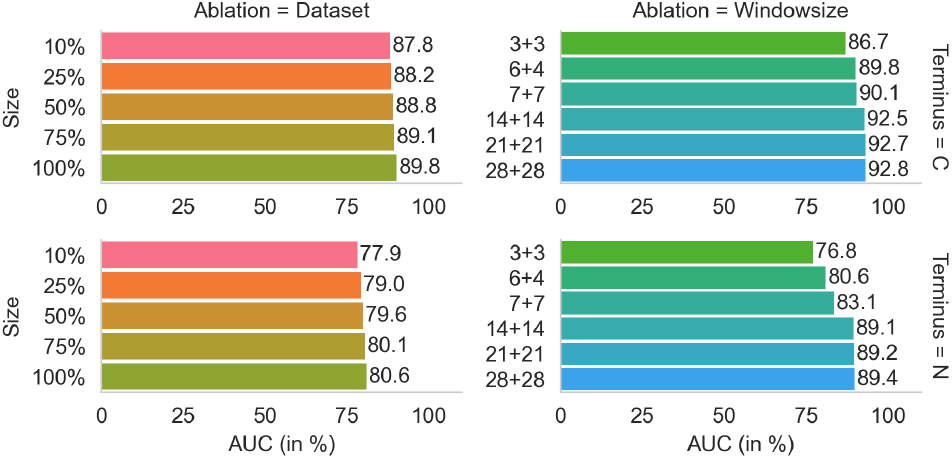
BiLSTM performance with smaller training datasets (left) and different window sizes (right) on C- and N-terminals (top and bottom rows).

#### Window size

As for window size, performance seemed to plateau when 14 residues were used at each side of the cleavage site, with an improvement of only around 0.3 p.p. when doubling the window size (Fig. 4 right, and Table 30 in Appendix A.5). Starting at a half-window size of 21, the BiLSTM models failed to learn anything, achieving no more than 54% training AUC. Introducing batch normalization layers [Ioffe and Szegedy, 2015] resolved the issue, and in this way the overall best AUC achieved with a window of 56 residues was 92.81% for C-terminals and 89.41% for N-terminals. Increasing the window size from 6 to 56 improved predictive performance by 12.61 pp. for N-terminals and only by 6.14 pp. for C-terminals.

#### Model size

To ensure that the performance numbers just mentioned were not limited by the capacity of the model, we tried to increase the number of units and layers by 50% and 100%, however, both models started to overfit before reaching the performance level of our base BiLSTM model. As we already applied 50% dropout [Srivastava et al., 2014], we progressively increased weight decay [Krogh and Hertz, 1991] and repeated training. In no scenario did any combination of weight decay and increased model size achieve the performance of our base BiLSTM model. For higher levels of weight decay, e.g., 1 × 10^*−*2^, training was severely limited and learning stopped at 87% AUC. Lower levels of weight decay, e.g., 1 × 10^*−*4^ were more competitive but still could not match the base BiLSTM performance, even in a setting with extended training epochs. The learning and generalization curves for the 50% size increase model were mirrored by those with 100% increased size. Exact results for each weight decay setting can be found in Table 31 in Appendix A.5.

### 4.2. Baseline results

Models re-trained on our dataset (PCM, Logistic regression, SVM, NetCleave, PUUPL, and Pepsickle) outperformed available pre-trained models (both versions of NetChop, PCPS and PCM). In all cases, our BiLSTM models outperform all to-date published methods and thus constitute a new state-of-the-art (Table 2). Interestingly, both PCM and the logistic regression classifier performed better than NetChop but worse than NetCleave. Both NetChop and NetCleave are based on MLPs, but while NetCleave was re-trained, NetChop was not, suggesting that retraining NetChop on our dataset would obtain higher performance than shown in the table. Importantly, such simple linear models achieved only three or four pp. fewer than the considerably more complicated BiLSTM. The fact that both linear models performed very competitively compared to the other deep architectures suggests that improvements in the training data were the major driver of the improved performance observed over the years. The SVM model performed slightly worse than logistic regression even though the hyperparameter optimization preferred a RBF kernel over the linear and polynomial kernels. The number of training samples was the bottleneck, and performance could be increased at great computational cost.

## 5. Conclusion

We benchmarked various deep learning architectures, including MLPs, CNNs, LSTMs, and Transformers, for the task of proteasomal cleavage prediction, and showed that several embedding techniques in combination with model architectures of vastly different scale and complexity all reach similar performance levels around 89.8% AUC for C-terminals and 80.6% AUC for N-terminals, while a simple linear model only achieves three or four p.p. fewer compared to such complicated models. Denoising techniques as well as trainable tokenizers appeared to offer limited to no, or even negative benefits, possibly because the MHC ligandomics dataset already covers most of the cleavage sites that can result in an MHC ligand. By increasing the size of the window around the cleavage site, we reached up to 92.8% and 89.4% AUC for C- and N-terminals, with evident plateaus in terms of training dataset and window size. Performance on the N-terminals dataset was more sensitive both to window size and the number of training samples, confirming the higher difficulty of predicting this process compared to C-terminus cleavage. When maximum predictive performance is desired, we suggest using our best-performing BiLSTM model with the largest possible window size that is feasible for the application at hand, while single-layer or multi-layer perceptrons can be used when increased efficiency is desired with only a slight penalty in predictive performance. The logistic regression classifier also presents a viable option when efficiency is the most important concern as its predictive performance is very competitive with all other deep neural network models.

Such saturated results suggest that different modeling choices of architectures, embeddings, or training regimes are unlikely to yield significantly better predictive performance in the future, although marginal gains could still be possible with even larger datasets. Based on this insight we speculate that further efforts to improve proteasomal cleavage prediction could focus on a more comprehensive modeling of the antigen pathway. For example, it has been shown that cleavage prediction benefits from additionally predicting the MHC binding affinity of the resulting peptides [Keşmir et al., 2002, Diez-Rivero et al., 2010]. Another possibility is that biological processes such as proteasomal cleavage are simply too noisy and random to allow more accurate predictions, in which case we may already be close to the boundary of what is possible to achieve *in silico*.

## Author contributions statement

I.Z. and B.M. conducted the experiments, analyzed the results, and wrote the manuscript. B.B. advised on the experimental setup and assisted in the interpretation of the results. E.D. conceived the experiments, prepared the dataset, advised on the experimental setup, assisted in the interpretation of the results, and wrote the manuscript. B.S. advised on the experimental setup, assisted in the interpretation of the results, and wrote the manuscript. All authors reviewed and approved the manuscript.

## Acknowledgments

E. D. was supported by the Helmholtz Association under the joint research school “Munich School for Data Science - MUDS” (Award Number HIDSS-0006). B. B. was supported by the Bavarian Ministry of Economic Affairs, Regional Development and Energy through the Center for Analytics – Data – Applications (ADA-Center) within the framework of BAYERN DIGITAL II (20-3410-2-9-8). B. B. was supported by the German Federal Ministry of Education and Research (BMBF) and the Munich Center for Machine Learning (MCML).

## A. Appendix

### A.1. Availability

As of October 2023, the publicly available proteasomal cleavage predictors listed in Table 2 are found at:

- ProteaSMM: The official website http://www.mhc-pathway.net/ does not reply. The model is reimplemented in the epytope framework: https://github.com/KohlbacherLab/epytope
- PCM is implemented in the epytope framework: https://github.com/KohlbacherLab/epytope
- NetChop 3.1: https://services.healthtech.dtu.dk/service.php?NetChop-3.1
- Pcleavage: http://crdd.osdd.net/raghava/pcleavage/
- PCPS: http://imed.med.ucm.es/Tools/pcps/
- NetCleave: https://github.com/pepamengual/NetCleave
- Pepsickle: https://github.com/pdxgx/pepsickle
- PUUPL: https://github.com/SchubertLab/proteasomal-cleavage-puupl
- This paper: https://github.com/ziegler-ingo/cleavage_benchmark

### A.2. Hyperparameters

**Table 3.** BiLSTM.

| Hyperparameter | Range<br>(Uniformly random choice) | Final value |  |
| --- | --- | --- | --- |
|  |  | C-terminal | N-terminal |
| Epochs trained | $\leq 25$ | 15 | 15 |
| Learning rate | $\{5 \times 10^{-5}, 10^{-4}, 3 \times 10^{-4}\}$ | $3 \times 10^{-4}$ | $3 \times 10^{-4}$ |
| Dropout rate | $\{0.45, 0.46, \dots, 0.51, 0.52\}$ | 0.5 | 0.5 |
| Linear layer size | [120, 181) | 164 | 179 |
| Embedding dimension | [50, 201) | 91 | 76 |
| LSTM size 1 | [220, 281) | 228 | 252 |
| LSTM size 2 | [450, 520) | 506 | 518 |

**Table 4.** BiLSTM+Attention.

| Hyperparameter | Range<br>(Uniformly random choice) | Final value |  |
| --- | --- | --- | --- |
|  |  | C-terminal | N-terminal |
| Epochs trained | $\leq 25$ | 20 | 20 |
| Learning rate | $\{3 \times 10^{-5}, 5 \times 10^{-5}, 8 \times 10^{-5}, 10^{-4}\}$ | $10^{-4}$ | $10^{-4}$ |
| Dropout rate | $\{0.45, 0.46, \dots, 0.51, 0.52\}$ | 0.5 | 0.5 |
| Linear layer size | [100, 181) | 147 | 150 |
| Embedding dimension | $\{120, 124, \dots, 216, 220\}$ | 216 | 216 |
| LSTM size | [64, 131) | 108 | 111 |
| Attention heads | {1, 2, 4} | 4 | 1 |

**Table 5.** BiLSTM+Prot2Vec.

| Hyperparameter | Range<br>(Uniformly random choice) | Final value |  |
| --- | --- | --- | --- |
|  |  | C-terminal | N-terminal |
| Epochs trained | $\leq 60$ | 60 | 60 |
| Learning rate | $\{8 \times 10^{-5}, 10^{-4}, 3 \times 10^{-4}, 5 \times 10^{-4}\}$ | $3 \times 10^{-4}$ | $3 \times 10^{-4}$ |
| Dropout rate | $\{0.45, 0.46, \dots, 0.51, 0.52\}$ | 0.5 | 0.5 |
| Linear layer size | [120, 180) | 145 | 139 |
| LSTM size | [480, 531) | 480 | 531 |

**Table 6.** CNN.

| Hyperparameter | Range<br>(Uniformly random choice) | Final value |  |
| --- | --- | --- | --- |
|  |  | C-terminal | N-terminal |
| Epochs trained | $\leq 60$ | 60 | 60 |
| Learning Rate | $\{8 \times 10^{-5}, 10^{-4}, 3 \times 10^{-4}, 5 \times 10^{-4}\}$ | $3 \times 10^{-4}$ | $3 \times 10^{-4}$ |
| Dropout Rate | $\{0, 0.02, \dots, 0.08, 0.1\}$ | 0.04 | 0.08 |
| Linear layer size 1 | [64, 101) | 79 | 89 |
| Linear layer size 2 | [15, 33) | 24 | 15 |
| Attention heads 1 | $\{1, 2, 3, 4, 5, 6\}$ | 3 | 4 |
| Attention heads 2 | $\{1, 2, 3, 4, 5, 6\}$ | 2 | 3 |
| Filter size 1 | - | 1 | 1 |
| Number filters 1 | [220, 301) | 220 | 249 |
| Number filters 2 | [220, 301) | 262 | 229 |
| Filter size 2a | $\{1, 3, 5, 7\}$ | 3 | 1 |
| Filter size 2b | $\{15, 17, 19, 21, 23, 25\}$ | 17 | 15 |
| Filter size 3c | $\{13, 15, 17, 19, 21, 23\}$ | 13 | 13 |
| Number filters 3 | [350, 431) | 398 | 400 |
| Filter size 3a | $\{11, 13, 15, 17, 19\}$ | 11 | 13 |
| Filter size 3b | $\{13, 15, 17, 19, 21, 23\}$ | 15 | 21 |
| Filter size 3c | $\{11, 13, 15, 17, 19\}$ | 19 | 15 |

**Table 7.** MLP.

| Hyperparameter | Range<br>(Uniformly random choice) | Final value |  |
| --- | --- | --- | --- |
|  |  | C-terminal | N-terminal |
| Epochs trained | $\leq 60$ | 30 | 30 |
| Learning rate | $\{10^{-4}, 5 \times 10^{-4}, 8 \times 10^{-4}, 10^{-3}\}$ | $10^{-3}$ | $10^{-3}$ |
| Dropout rate | $\{0.1, 0.12, \dots, 0.24, 0.26\}$ | 0.24 | 0.24 |
| Linear layer size | [120, 201) | 144 | 167 |

**Table 8.** Pepsickle.

| Table 3: Hyperparameters |  |  |  |  |
| --- | --- | --- | --- | --- |
| Hyperparameter |  | Range<br>(Uniformly random choice) | Final value |  |
|  |  |  | C-terminal | N-terminal |
| Sequence-Net | Epochs trained | $\leq 100$ | 20 | 10 |
| | Learning rate | $10^{-3}$ | $10^{-3}$ | $10^{-3}$ |
| | Dropout rate | $\{0.1, 0.12, \dots, 0.34, 0.36\}$ | 0.34 | 0.32 |
| | Linear layer size 1 | $[120, 201)$ | 187 | 175 |
| | Linear layer size 2 | $[60, 141)$ | 71 | 96 |
| | Linear layer size 3 | $[20, 81)$ | 20 | 44 |
| Motif-Net | Epochs trained | $\leq 100$ | 20 | 10 |
| | Learning rate | $\{10^{-3}\}$ | $10^{-3}$ | $10^{-3}$ |
| | Dropout rate | $\{0.1, 0.12, \dots, 0.24, 0.26\}$ | 0.12 | 0.12 |
| | Linear layer size 1 | $[120, 201)$ | 200 | 200 |
| | Linear layer size 2 | $[20, 101)$ | 23 | 23 |

**Table 9.**
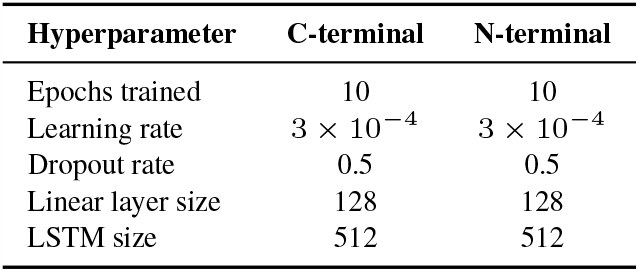
BiLSTM+ESM2.

| Hyperparameter | C-terminal | N-terminal |
| --- | --- | --- |
| Epochs trained | 10 | 10 |
| Learning rate | $3 \times 10^{-4}$ | $3 \times 10^{-4}$ |
| Dropout rate | 0.5 | 0.5 |
| Linear layer size | 128 | 128 |
| LSTM size | 512 | 512 |

**Table 10.** ESM2.

| Hyperparameter | C-terminal | N-terminal |
| --- | --- | --- |
| Epochs trained | 3 | 3 |
| Learning rate | $2 \times 10^{-5}$ | $2 \times 10^{-5}$ |
| Dropout rate | 0.5 | 0.5 |

**Table 11.**
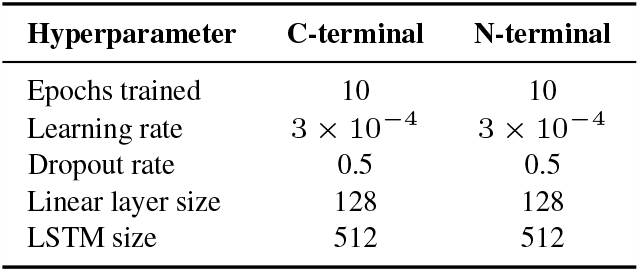
BiLSTM+T5.

**Table 12.** BiLSTM+BBPE1, BiLSTM+BBPE50, BiLSTM+WP50.

| Hyperparameter | C-terminal | N-terminal |
| --- | --- | --- |
| Epochs trained | 15 | 15 |
| Learning rate | $10^{-4}$ | $10^{-4}$ |
| Dropout rate | 0.5 | 0.5 |
| Embedding dimension | 150 | 150 |
| Linear layer size | 128 | 128 |
| LSTM size | 512 | 512 |

**Table 13.** BiLSTM+FwBw.

| Hyperparameter | C-terminal | N-terminal |
| --- | --- | --- |
| Epochs trained | 15 | 15 |
| Learning rate | $10^{-4}$ | $10^{-4}$ |
| Dropout rate | 0.5 | 0.5 |
| Linear layer size 1 | 128 | 128 |
| LSTM size 1 | 128 | 128 |
| LSTM size 2 | 512 | 512 |
| Sequence encoding embedding dimension | 100 | 100 |
| Sequence encoding BiLSTM size | 200 | 200 |

**Table 14.** Logistic Regression.

| Hyperparameter | C-terminal | N-terminal |
| --- | --- | --- |
| Maximum Iteration | 100 | 100 |
| Cs | 10 | 10 |
| Penalty | L2 | L2 |
| Fit Intercept | Yes | Yes |
| Intercept Scaling | 1 | 1 |
| Solver | LBFGS | LBFGS |
| Tolerance | $10^{-4}$ | $10^{-4}$ |

**PUUPL:** See configuration files at https://github.com/SchubertLab/proteasomal-cleavage-puupl/tree/main/configs

**Table 15.** Denoising methods (Co-Teaching, Co-Teaching+, JoCoR, DivideMix, Noise adaptation layer)

| Hyperparameter | Co-teaching | Co-teaching+ | JoCoR | DivideMix | Noise adaptation layer |
| --- | --- | --- | --- | --- | --- |
| Number scale-up epochs | 10 | 10 | 10 | 5 | - |
| Noise rate | 0.2 | 0.2 | 0.2 | - | - |
| Forget rate | 0.2 | 0.2 | 0.1 | - | - |
| Exponent | 1 | 1 | 1 | - | - |
| Number warm-up epochs | - | - | - | 1 | 1 |
| $\alpha$ | - | - | - | 0.5 | - |
| $\lambda_u$ | - | - | - | 0 | - |
| Probability threshold | - | - | - | 0.5 | - |
| Temperature | - | - | - | 0.5 | - |
| $\beta$ | - | - | - | - | 0.8 |

### A.3. Results without denoising methods

**Table 16.** Model performances without denoising.

| Terminal | Prio. | Models | AUC | ACC | s/epoch | Epochs | Parameters | Trainable |
| --- | --- | --- | --- | --- | --- | --- | --- | --- |
| C-terminal | 1 | BiLSTM | 89.85 ± 0.11 | 87.66 ± 0.07 | 25 | 15 | 4 655 984 | 4 655 984 |
| | | BiLSTM+Attention | 88.90 ± 0.11 | 87.09 ± 0.09 | 20 | 20 | 1 632 391 | 1 632 391 |
| | | BiLSTM+Prot2Vec | 88.80 ± 0.04 | 87.26 ± 0.06 | 20 | 15 | 16 009 371 | 5 830 371 |
| | | CNN | 87.77 ± 0.14 | 86.25 ± 0.13 | 45 | 60 | 16 084 198 | 16 084 198 |
| | | MLP | 88.86 ± 0.11 | 87.22 ± 0.08 | 4 | 30 | 30 529 | 30 529 |
| | 2 | BiLSTM+ESM2 | 88.53 ± 0.07 | 87.01 ± 0.03 | 330 | 10 | 152 998 267 | 4 858 113 |
| | | ESM2 | 88.74 ± 0.13 | 86.37 ± 1.46 | 900 | 3 | 148 140 188 | 148 140 188 |
| | | BiLSTM+T5 | 86.70 ± 0.05 | 85.91 ± 0.14 | 780 | 10 | 1 214 572 801 | 6 430 977 |
| | | BiLSTM+BBPE1 | 85.91 ± 0.11 | 85.83 ± 0.10 | 13 | 15 | 5 319 409 | 5 319 409 |
| | | BiLSTM+BBPE50 | 69.14 ± 0.17 | 82.41 ± 0.11 | 12 | 15 | 12 669 409 | 12 669 409 |
| | | BiLSTM+WP50 | 73.95 ± 0.24 | 82.99 ± 0.08 | 16 | 15 | 12 669 409 | 12 669 409 |
| | | BiLSTM+FwBw | 87.57 ± 0.07 | 86.75 ± 0.09 | 120 | 15 | 4 315 369 | 4 315 369 |
| N-terminal | 1 | BiLSTM | 80.62 ± 0.11 | 83.82 ± 0.12 | 25 | 15 | 5 096 135 | 5 096 135 |
| | | BiLSTM+Attention | 79.92 ± 0.10 | 83.63 ± 0.13 | 20 | 20 | 1 718 233 | 1 718 233 |
| | | BiLSTM+Prot2Vec | 80.31 ± 0.09 | 83.81 ± 0.07 | 20 | 15 | 16 772 049 | 6 593 049 |
| | | CNN | 77.91 ± 1.17 | 82.95 ± 0.26 | 45 | 60 | 15 237 057 | 15 237 057 |
| | | MLP | 80.46 ± 0.10 | 83.76 ± 0.13 | 4 | 30 | 35 405 | 35 405 |
| | 2 | BiLSTM+ESM2 | 79.59 ± 0.12 | 83.55 ± 0.07 | 360 | 10 | 152 998 267 | 4 858 113 |
| | | ESM2 | 79.96 ± 0.14 | 82.52 ± 0.57 | 900 | 3 | 148 140 188 | 148 140 188 |
| | | BiLSTM+T5 | 78.48 ± 0.15 | 83.06 ± 0.18 | 780 | 10 | 1 214 572 801 | 6 430 977 |
| | | BiLSTM+BBPE1 | 76.12 ± 0.06 | 82.61 ± 0.07 | 13 | 15 | 5 319 409 | 5 319 409 |
| | | BiLSTM+BBPE50 | 69.42 ± 0.20 | 82.15 ± 0.11 | 12 | 15 | 12 669 409 | 12 669 409 |
| | | BiLSTM+WP50 | 69.33 ± 0.10 | 81.91 ± 0.14 | 15 | 15 | 12 669 409 | 12 669 409 |
| | | BiLSTM+FwBw | 78.75 ± 0.13 | 82.23 ± 0.07 | 120 | 15 | 4 315 369 | 4 315 369 |

### A.4. Results with denoising methods

**Table 17.** BiLSTM with denoising.

| Terminal | Denoising methods | AUC | ACC | s/epoch | Epochs | Parameters | Trainable |
| --- | --- | --- | --- | --- | --- | --- | --- |
| C-terminal | Co-Teaching | 87.75 ± 0.09 | 86.80 ± 0.06 | 41 | 15 | 4 655 984 | 4 655 984 |
| | Co-Teaching+ | 87.75 ± 0.09 | 86.80 ± 0.06 | 41 | 15 | 4 655 984 | 4 655 984 |
| | JoCoR | 84.70 ± 0.22 | 85.36 ± 0.06 | 41 | 15 | 4 655 984 | 4 655 984 |
|  | DivideMix | 86.25 | 84.02 | 210 | 15 | 4 656 149 | 4 656 149 |
| | Noise Adaptation Layer | 89.11 ± 0.06 | 87.36 ± 0.04 | 22 | 15 | 4 656 149 | 4 656 149 |
| N-terminal | Co-Teaching | 78.74 ± 0.11 | 83.41 ± 0.11 | 44 | 15 | 5 096 135 | 5 096 135 |
| | Co-Teaching+ | 78.74 ± 0.11 | 83.41 ± 0.11 | 44 | 15 | 5 096 135 | 5 096 135 |
| | JoCoR | 74.09 ± 0.25 | 82.83 ± 0.14 | 44 | 15 | 5 096 135 | 5 096 135 |
|  | DivideMix | 77.08 | 81.52 | 210 | 15 | 5 096 315 | 5 096 315 |
| | Noise Adaptation Layer | 70.60 ± 0.10 | 83.81 ± 0.07 | 23 | 15 | 5 096 315 | 5 096 315 |

**Table 18.** BiLSTM+Attention with denoising.

| Terminal | Denoising methods | AUC | ACC | s/epoch | Epochs | Parameters | Trainable |
| --- | --- | --- | --- | --- | --- | --- | --- |
| C-terminal | Co-Teaching | 87.10 ± 0.05 | 86.42 ± 0.10 | 40 | 20 | 1 632 391 | 1 632 391 |
| | Co-Teaching+ | 87.10 ± 0.05 | 86.42 ± 0.10 | 40 | 20 | 1 632 391 | 1 632 391 |
| | JoCoR | 82.17 ± 0.29 | 85.46 ± 0.06 | 38 | 20 | 1 632 391 | 1 632 391 |
| | Noise Adaptation Layer | 88.93 ± 0.06 | 87.24 ± 0.08 | 18 | 20 | 1 632 539 | 1 632 539 |
| N-terminal | Co-Teaching | 77.96 ± 0.11 | 83.19 ± 0.07 | 43 | 20 | 1 718 233 | 1 718 233 |
| | Co-Teaching+ | 77.96 ± 0.11 | 83.19 ± 0.07 | 42 | 20 | 1 718 233 | 1 718 233 |
| | JoCoR | 70.16 ± 0.70 | 82.12 ± 0.14 | 41 | 20 | 1 718 233 | 1 718 233 |
| | Noise Adaptation Layer | 78.89 ± 0.11 | 83.59 ± 0.06 | 22 | 20 | 1 718 384 | 1 718 384 |

**Table 19.** BiLSTM+Prot2Vec with denoising.

| Terminal | Denoising methods | AUC | ACC | s/epoch | Epochs | Parameters | Trainable |
| --- | --- | --- | --- | --- | --- | --- | --- |
| C-terminal | Co-Teaching | 86.94 ± 0.12 | 87.30 ± 0.07 | 32 | 15 | 16 009 371 | 5 830 371 |
| | Co-Teaching+ | 86.94 ± 0.12 | 87.30 ± 0.07 | 31 | 15 | 16 009 371 | 5 830 371 |
| | JoCoR | 76.18 ± 0.16 | 82.31 ± 0.12 | 32 | 15 | 16 009 371 | 5 830 371 |
| | Noise Adaptation Layer | 88.81 ± 0.08 | 87.21 ± 0.06 | 15 | 15 | 16 009 517 | 5 830 517 |
| N-terminal | Co-Teaching | 78.32 ± 0.12 | 83.56 ± 0.07 | 40 | 15 | 16 772 049 | 6 593 049 |
| | Co-Teaching+ | 78.32 ± 0.12 | 83.56 ± 0.07 | 40 | 15 | 16 772 049 | 6 593 049 |
| | JoCoR | 63.29 ± 1.08 | 81.83 ± 0.12 | 40 | 15 | 16 772 049 | 6 593 049 |
| | Noise Adaptation Layer | 80.56 ± 0.15 | 83.71 ± 0.08 | 19 | 15 | 16 772 189 | 6 593 189 |

**Table 20.**
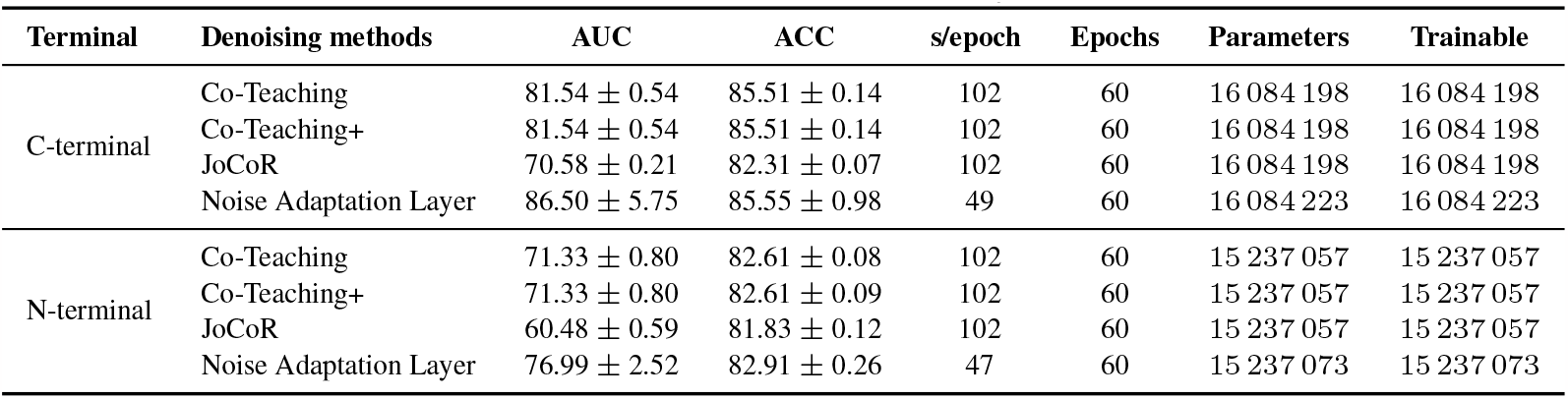
CNN with denoising.

**Table 21.** MLP with denoising.

| Terminal | Denoising methods | AUC | ACC | s/epoch | Epochs | Parameters | Trainable |
| --- | --- | --- | --- | --- | --- | --- | --- |
| C-terminal | Co-Teaching | 87.37 ± 0.10 | 86.72 ± 0.06 | 6 | 30 | 30 529 | 30 529 |
| | Co-Teaching+ | 87.37 ± 0.10 | 86.72 ± 0.06 | 6 | 30 | 30 529 | 30 529 |
| | JoCoR | 85.99 ± 0.09 | 85.63 ± 0.04 | 5 | 30 | 30 529 | 30 529 |
| | Noise Adaptation Layer | 88.70 ± 0.09 | 87.19 ± 0.07 | 3 | 30 | 30 674 | 30 674 |
| N-terminal | Co-Teaching | 78.30 ± 0.06 | 83.38 ± 0.10 | 6 | 30 | 35 405 | 35 405 |
| | Co-Teaching+ | 78.30 ± 0.06 | 83.38 ± 0.10 | 5 | 30 | 35 405 | 35 405 |
| | JoCoR | 76.21 ± 0.05 | 81.97 ± 0.14 | 5 | 30 | 35 405 | 35 405 |
| | Noise Adaptation Layer | 79.88 ± 0.10 | 83.65 ± 0.10 | 4 | 30 | 35 573 | 35 573 |

**Table 22.** BiLSTM+ESM2 with denoising.

| Terminal | Denoising methods | AUC | ACC | s/epoch | Epochs | Parameters | Trainable |
| --- | --- | --- | --- | --- | --- | --- | --- |
| C-terminal | Co-Teaching | 87.37 ± 0.02 | 86.65 ± 0.05 | 660 | 10 | 152 998 267 | 4 858 113 |
| | Co-Teaching+ | 87.37 ± 0.02 | 86.65 ± 0.05 | 660 | 10 | 152 998 267 | 4 858 113 |
| | JoCoR | 84.00 ± 0.11 | 84.82 ± 0.08 | 660 | 10 | 152 998 267 | 4 858 113 |
| | Noise Adaptation Layer | 88.55 ± 0.08 | 87.00 ± 0.06 | 330 | 10 | 152 998 396 | 4 858 242 |
| N-terminal | Co-Teaching | 78.30 ± 0.08 | 83.42 ± 0.08 | 660 | 10 | 152 998 267 | 4 858 113 |
| | Co-Teaching+ | 78.30 ± 0.08 | 83.42 ± 0.08 | 660 | 10 | 152 998 267 | 4 858 113 |
| | JoCoR | 71.41 ± 0.36 | 81.86 ± 0.13 | 660 | 10 | 152 998 267 | 4 858 113 |
| | Noise Adaptation Layer | 79.51 ± 0.07 | 83.50 ± 0.11 | 360 | 10 | 152 998 396 | 4 858 242 |

**Table 23.** ESM2 with denoising.

| Terminal | Denoising methods | AUC | ACC | s/epoch | Epochs | Parameters | Trainable |
| --- | --- | --- | --- | --- | --- | --- | --- |
| C-terminal | Co-Teaching | 87.99 ± 0.18 | 82.55 ± 0.83 | 2160 | 3 | 148 140 188 | 148 140 188 |
| | Co-Teaching+ | 87.99 ± 0.18 | 85.55 ± 0.83 | 2100 | 3 | 148 140 188 | 148 140 188 |
| | JoCoR | 87.12 ± 0.22 | 86.29 ± 0.84 | 2100 | 3 | 148 140 188 | 148 140 188 |
| | Noise Adaptation Layer | 88.68 ± 1.90 | 77.71 ± 11.67 | 960 | 3 | 148 140 222 | 148 140 222 |
| N-terminal | Co-Teaching | 78.90 ± 1.03 | 81.83 ± 2.59 | 2220 | 3 | 148 140 188 | 148 140 188 |
| | Co-Teaching+ | 78.90 ± 1.03 | 81.83 ± 2.59 | 2220 | 3 | 148 140 188 | 148 140 188 |
| | JoCoR | 77.17 ± 0.84 | 82.59 ± 0.28 | 2160 | 3 | 148 140 188 | 148 140 188 |
| | Noise Adaptation Layer | 79.82 ± 0.48 | 70.27 ± 14.85 | 930 | 3 | 148 140 222 | 148 140 222 |

**Table 24.** BiLSTM+T5 with denoising.

| Terminal | Denoising methods | AUC | ACC | s/epoch | Epochs | Parameters | Trainable |
| --- | --- | --- | --- | --- | --- | --- | --- |
| C-terminal | Co-Teaching | 86.41 ± 0.20 | 86.31 ± 0.17 | 1500 | 10 | 1 214 572 801 | 6 430 977 |
| | Co-Teaching+ | 86.41 ± 0.20 | 86.31 ± 0.17 | 1500 | 10 | 1 214 572 801 | 6 430 977 |
| | JoCoR | 83.72 ± 0.15 | 84.01 ± 0.06 | 1500 | 10 | 1 214 572 801 | 6 430 977 |
| | Noise Adaptation Layer | 87.21 ± 0.17 | 86.12 ± 0.30 | 720 | 10 | 1 214 572 930 | 6 431 106 |
| N-terminal | Co-Teaching | 77.28 ± 0.11 | 82.84 ± 0.10 | 1560 | 10 | 1 214 572 801 | 6 430 977 |
| | Co-Teaching+ | 77.28 ± 0.11 | 85.91 ± 0.14 | 1560 | 10 | 1 214 572 801 | 6 430 977 |
| | JoCoR | 68.79 ± 0.17 | 81.83 ± 0.12 | 1500 | 10 | 1 214 572 801 | 6 430 977 |
| | Noise Adaptation Layer | 78.30 ± 0.05 | 82.82 ± 0.18 | 780 | 10 | 1 214 572 930 | 6 431 106 |

**Table 25.** BiLSTM+BBPE1 with denoising.

| Terminal | Denoising methods | AUC | ACC | s/epoch | Epochs | Parameters | Trainable |
| --- | --- | --- | --- | --- | --- | --- | --- |
| C-terminal | Co-Teaching | 82.48 ± 0.12 | 85.42 ± 0.06 | 33 | 15 | 5 319 409 | 5 319 409 |
| | Co-Teaching+ | 82.48 ± 0.12 | 85.42 ± 0.06 | 34 | 15 | 5 319 409 | 5 319 409 |
| | JoCoR | 60.39 ± 0.37 | 82.20 ± 0.07 | 33 | 15 | 5 319 409 | 5 319 409 |
| | Noise Adaptation Layer | 85.70 ± 0.21 | 85.72 ± 0.20 | 17 | 15 | 5 319 538 | 5 319 538 |
| N-terminal | Co-Teaching | 74.50 ± 0.27 | 82.59 ± 0.09 | 34 | 15 | 5 319 409 | 5 319 409 |
| | Co-Teaching+ | 74.50 ± 0.27 | 82.59 ± 0.09 | 34 | 15 | 5 319 409 | 5 319 409 |
| | JoCoR | 63.11 ± 0.36 | 81.88 ± 0.12 | 33 | 15 | 5 319 409 | 5 319 409 |
| | Noise Adaptation Layer | 76.08 ± 0.14 | 82.53 ± 0.09 | 17 | 15 | 5 319 538 | 5 319 538 |

**Table 26.** BiLSTM+BBPE50 with denoising.

| Terminal | Denoising methods | AUC | ACC | s/epoch | Epochs | Parameters | Trainable |
| --- | --- | --- | --- | --- | --- | --- | --- |
| C-terminal | Co-Teaching | 66.44 ± 0.18 | 82.28 ± 0.08 | 30 | 15 | 12 669 409 | 12 669 409 |
| | Co-Teaching+ | 66.44 ± 0.18 | 82.28 ± 0.08 | 30 | 15 | 12 669 409 | 12 669 409 |
| | JoCoR | 53.20 ± 0.22 | 82.20 ± 0.07 | 30 | 15 | 12 669 409 | 12 669 409 |
| | Noise Adaptation Layer | 68.67 ± 0.13 | 82.38 ± 0.13 | 16 | 15 | 12 669 538 | 12 669 538 |
| N-terminal | Co-Teaching | 66.49 ± 0.23 | 82.05 ± 0.13 | 31 | 15 | 12 669 409 | 12 669 409 |
| | Co-Teaching+ | 66.49 ± 0.23 | 82.05 ± 0.13 | 32 | 15 | 12 669 409 | 12 669 409 |
| | JoCoR | 58.99 ± 0.81 | 81.88 ± 0.12 | 31 | 15 | 12 669 409 | 12 669 409 |
| | Noise Adaptation Layer | 68.88 ± 0.37 | 82.14 ± 0.14 | 15 | 15 | 12 669 538 | 12 669 538 |

**Table 27.**
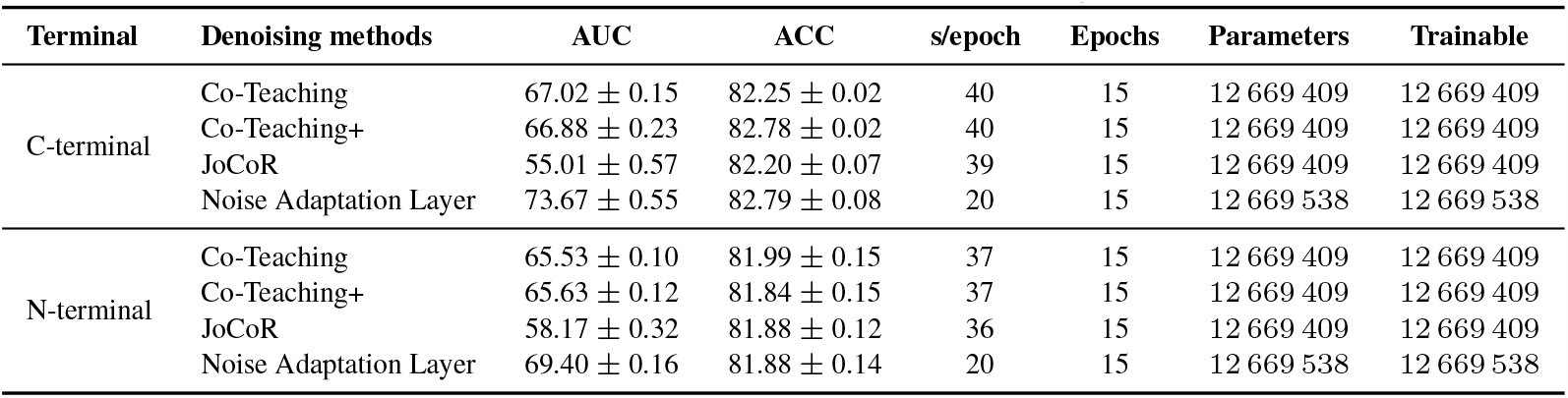
BiLSTM+WP50 with denoising.

**Table 28.**
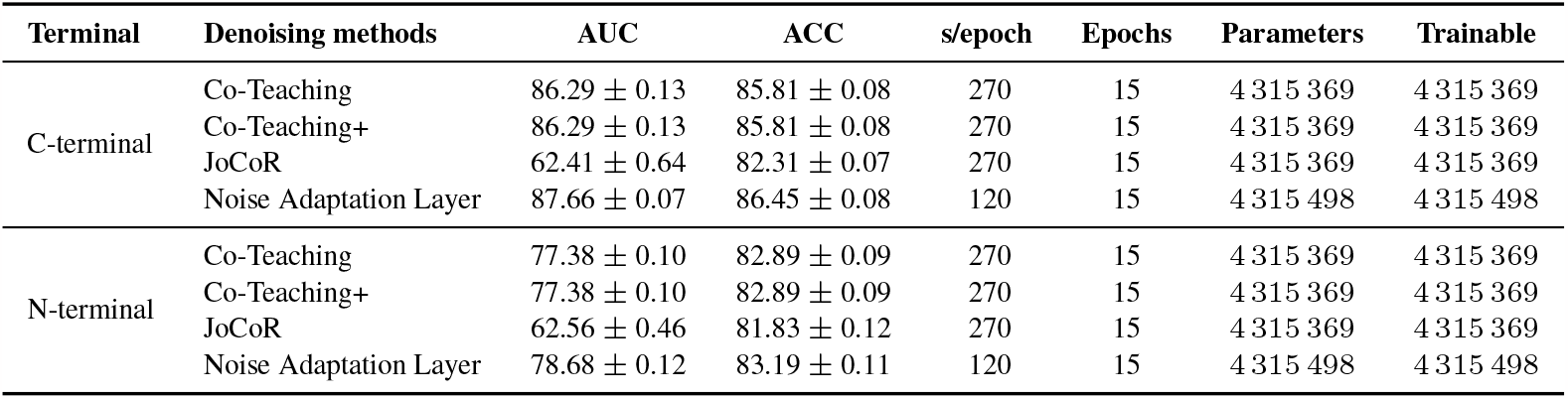
BiLSTM+FwBw with denoising.

| Terminal | Denoising methods | AUC | ACC | s/epoch | Epochs | Parameters | Trainable |
| --- | --- | --- | --- | --- | --- | --- | --- |
| C-terminal | Co-Teaching | 86.29 ± 0.13 | 85.81 ± 0.08 | 270 | 15 | 4 315 369 | 4 315 369 |
| | Co-Teaching+ | 86.29 ± 0.13 | 85.81 ± 0.08 | 270 | 15 | 4 315 369 | 4 315 369 |
| | JoCoR | 62.41 ± 0.64 | 82.31 ± 0.07 | 270 | 15 | 4 315 369 | 4 315 369 |
| | Noise Adaptation Layer | 87.66 ± 0.07 | 86.45 ± 0.08 | 120 | 15 | 4 315 498 | 4 315 498 |
| N-terminal | Co-Teaching | 77.38 ± 0.10 | 82.89 ± 0.09 | 270 | 15 | 4 315 369 | 4 315 369 |
| | Co-Teaching+ | 77.38 ± 0.10 | 82.89 ± 0.09 | 270 | 15 | 4 315 369 | 4 315 369 |
| | JoCoR | 62.56 ± 0.46 | 81.83 ± 0.12 | 270 | 15 | 4 315 369 | 4 315 369 |
| | Noise Adaptation Layer | 78.68 ± 0.12 | 83.19 ± 0.11 | 120 | 15 | 4 315 498 | 4 315 498 |

### A.5. Ablation results

**Table 29.**
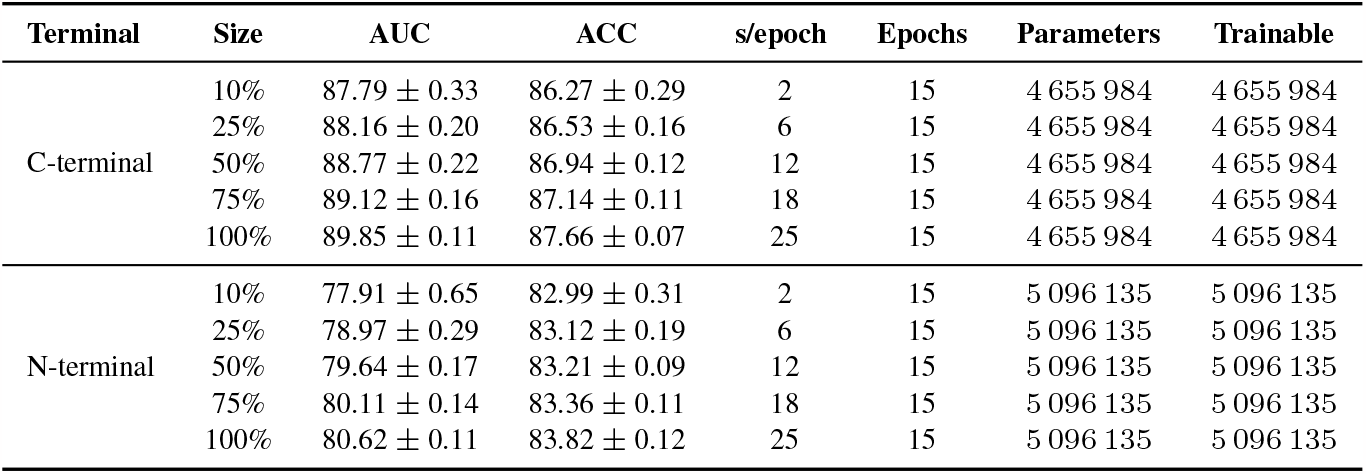
Dataset size ablation of BiLSTM.

| Terminal | Size | AUC | ACC | s/epoch | Epochs | Parameters | Trainable |
| --- | --- | --- | --- | --- | --- | --- | --- |
| C-terminal | 10% | 87.79 ± 0.33 | 86.27 ± 0.29 | 2 | 15 | 4 655 984 | 4 655 984 |
| | 25% | 88.16 ± 0.20 | 86.53 ± 0.16 | 6 | 15 | 4 655 984 | 4 655 984 |
| | 50% | 88.77 ± 0.22 | 86.94 ± 0.12 | 12 | 15 | 4 655 984 | 4 655 984 |
| | 75% | 89.12 ± 0.16 | 87.14 ± 0.11 | 18 | 15 | 4 655 984 | 4 655 984 |
| | 100% | 89.85 ± 0.11 | 87.66 ± 0.07 | 25 | 15 | 4 655 984 | 4 655 984 |
| N-terminal | 10% | 77.91 ± 0.65 | 82.99 ± 0.31 | 2 | 15 | 5 096 135 | 5 096 135 |
| | 25% | 78.97 ± 0.29 | 83.12 ± 0.19 | 6 | 15 | 5 096 135 | 5 096 135 |
| | 50% | 79.64 ± 0.17 | 83.21 ± 0.09 | 12 | 15 | 5 096 135 | 5 096 135 |
| | 75% | 80.11 ± 0.14 | 83.36 ± 0.11 | 18 | 15 | 5 096 135 | 5 096 135 |
| | 100% | 80.62 ± 0.11 | 83.82 ± 0.12 | 25 | 15 | 5 096 135 | 5 096 135 |

**Table 30.** Window size ablation of BiLSTM.

| Terminal | Halfwindow Size | AUC | ACC | s/epoch | Epochs | Parameters | Trainable |
| --- | --- | --- | --- | --- | --- | --- | --- |
| C-terminal | 3 | 86.67 ± 0.12 | 85.62 ± 0.10 | 17 | 15 | 4 655 984 | 4 655 984 |
| | 7 | 90.07 ± 0.12 | 87.94 ± 0.11 | 30 | 15 | 4 655 984 | 4 655 984 |
| | 14 | 92.50 ± 0.09 | 89.71 ± 0.07 | 57 | 15 | 4 655 984 | 4 655 984 |
| | 21 | 92.74 ± 0.11 | 89.90 ± 0.09 | 84 | 15 | 4 656 648 | 4 656 648 |
| | 28 | 92.81 ± 0.17 | 90.01 ± 0.16 | 109 | 15 | 4 656 648 | 4 656 648 |
| N-terminal | 3 | 76.80 ± 0.24 | 82.76 ± 0.05 | 21 | 15 | 5 096 135 | 5 096 135 |
| | 7 | 83.12 ± 0.18 | 84.82 ± 0.09 | 34 | 15 | 5 096 135 | 5 096 135 |
| | 14 | 89.13 ± 0.11 | 88.17 ± 0.13 | 64 | 15 | 5 096 135 | 5 096 135 |
| | 21 | 89.24 ± 0.22 | 88.53 ± 0.11 | 90 | 15 | 5 096 829 | 5 096 829 |
| | 28 | 89.41 ± 0.11 | 88.78 ± 0.14 | 115 | 15 | 5 096 829 | 5 096 829 |

**Table 31.** Size variations with weight decay for BiLSTM on C-terminal with a halfwindow size of 28.

| Modal size | Weight decay | Max AUC | Epoch | s/epoch | Epochs trained | Parameters | Trainable |
| --- | --- | --- | --- | --- | --- | --- | --- |
| 50% | 0 | 91.89 | 5 | 210 | 15 | 10 464 925 | 10 464 925 |
| | $1 \times 10^{-2}$ | 87.25 | 15 | 210 | 15 | 10 464 925 | 10 464 925 |
| | $1 \times 10^{-3}$ | 89.55 | 15 | 210 | 15 | 10 464 925 | 10 464 925 |
| | $1 \times 10^{-4}$ | 91.68 | 19 | 210 | 20 | 10 464 925 | 10 464 925 |
| | $1 \times 10^{-5}$ | 91.85 | 6 | 210 | 15 | 10 464 925 | 10 464 925 |
| 100% | 0 | 91.84 | 4 | 330 | 15 | 10 464 925 | 10 464 925 |
| | $1 \times 10^{-2}$ | 87.31 | 15 | 330 | 15 | 10 464 925 | 10 464 925 |
| | $1 \times 10^{-3}$ | 89.72 | 15 | 330 | 15 | 10 464 925 | 10 464 925 |
| | $1 \times 10^{-4}$ | 91.70 | 18 | 330 | 20 | 10 464 925 | 10 464 925 |
| | $1 \times 10^{-5}$ | 91.77 | 5 | 330 | 15 | 10 464 925 | 10 464 925 |

